# High-efficiency optogenetic silencing with soma-targeted anion-conducting channelrhodopsins

**DOI:** 10.1101/225847

**Authors:** Mathias Mahn, Lihi Gibor, Katayun Cohen-Kashi Malina, Pritish Patil, Yoav Printz, Shir Oring, Rivka Levy, Ilan Lampl, Ofer Yizhar

## Abstract

Optogenetic silencing allows time-resolved functional interrogation of defined neuronal populations. However, the limitations of inhibitory optogenetic tools impose stringent constraints on experimental paradigms. The high light power requirement of light-driven ion pumps and their effects on intracellular ion homeostasis pose unique challenges, particularly in experiments that demand inhibition of a widespread neuronal population *in vivo. Guillardia theta* anion-conducting channelrhodopsins (GtACRs) are promising in this regard, due to their high single-channel conductance and favorable photon-ion stoichiometry. However, GtACRs show poor membrane targeting in mammalian cells, and the activity of such channels can cause transient excitation in the axon due to an excitatory chloride reversal potential in this compartment. Here we address both problems by enhancing membrane targeting and subcellular compartmentalization of GtACRs. The resulting GtACR-based optogenetic tools show improved photocurrents, greatly reduced axonal excitation, high light sensitivity and rapid kinetics, allowing highly efficient inhibition of neuronal activity in the mammalian brain.

## Introduction

Perturbation of neuronal activity is a fundamental aspect of neuroscience research, often used to gain insight into the functional roles of particular brain regions, circuits and cell types^1^. Optogenetic tools have greatly enhanced the precision with which such manipulations can be performed^2^, providing both temporal precision and cell-type specificity to experiments aimed at defining the roles of individual neural circuit components in neural computation or animal behavior. Current optogenetic approaches for silencing of neurons are mainly based on the light-activated microbial rhodopsins halorhodopsin^3,4^, archaerhodopsin^5^ and cruxhalorhodopsin^6^. These proteins pump ions across the neuronal membrane with millisecond kinetics, independently of the electrochemical gradient. These tools allow neuronal silencing with precise temporal onset and offset^5,7,8^. However, ion-pumping rhodopsins possess several characteristics that impose substantial constraints on the experimental paradigm and complicate the interpretation of experimental outcomes. These limitations become even more pronounced in cases where neuronal silencing is required for extended periods of time. The unfavorable stoichiometry of one transported ion for each absorbed photon necessitates continuous illumination at high light power. The resulting tissue heating^9,10^ and phototoxicity^11^ restrict the optically addressable brain volume that can be efficiently silenced. Furthermore, ion-pumping microbial rhodopsins exhibit a decline in photocurrent amplitudes of up to 90% within a minute of illumination, leading to reduced silencing efficacy over time ^12,8,13^. Because of their insensitivity to electrochemical gradients, ion-pumping microbial rhodopsins can shift the concentrations of intracellular ions to non-physiological levels. In the case of halorhodopsin, this can lead to accumulation of chloride in the neuron, inducing changes in the reversal potential of GABAergic synapses ^14^. While in the case of archaerhodopsin this can increase the intracellular pH, inducing action potential-independent Ca^2+^ influx and elevated spontaneous vesicle release ^13^. Furthermore, the hyperpolarization mediated by ion-pumping activity together with the fast off kinetics can lead to an increased firing rate upon termination of the illumination ^6,13^.

Anion-conducting channelrhodopsins (ACRs), a newly established set of optogenetic tools^15,16,17^, are distinct from ion-pumping rhodopsins in two major aspects: first, they can conduct multiple ions during each photoreaction cycle. This increased photocurrent yield per photon makes channelrhodopsins superior in terms of their operational light-sensitivity. Second, conducting ions according to the reversal potential, ACRs are more likely to avoid non-physiological changes in ion concentration gradients. A light-gated chloride conductance will shunt membrane depolarization, which can be used to effectively clamp the neuronal membrane potential to the reversal potential of chloride, given that the ion permeability is sufficiently high. Anion-conducting channelrhodopsins could therefore relieve constrains imposed by ion-pumping rhodopsins. The naturally-occurring anion-conducting channelrhodopsins (nACRs) from the cryptophyte alga *Guillardia theta* ^16^ are particularly interesting in this regard. These channelrhodopsins, named GtACR1 and GtACR2, have near-perfect anion selectivity and produce large photocurrents in mammalian cells, owing to a higher single-channel conductance than that of the known cation-conducting channelrhodopsins^16,18^. While GtACRs were shown to inhibit behavior in the fruit fly^19,20^ and larval zebrafish ^21^, they have not yet been applied to mammalian systems, most likely due to poor membrane targeting and complex activity in the axonal compartment. To overcome these limitations of GtACRs and thereby of optogenetic inhibition in general, we generated several membrane targeting-enhanced GtACR variants, converging onto soma-targeted GtACR2 (stGtACR2), a fusion construct that combines GtACR2 with a C-terminal targeting motif from the soma-localized potassium channel Kv2.1^22^. We demonstrate here that stGtACR2 shows increased membrane targeting, extremely high anion photocurrents and reduced axonal excitation, making it the most effective tool for optogenetic inhibition at the cell soma to date.

## Results

### GtACR2 efficiently silences neuronal activity *in vitro* and *in vivo*

To determine the utility of ACRs for silencing of neurons we first expressed the three previously-described blue light-activated ACRs, GtACR2 ^16^, iC++ ^17^, and iChloC ^15^, in cultured rat hippocampal neurons by adeno associated virus (AAV)-mediated gene transfer. Whole-cell patch-clamp recordings from GtACR2-expressing neurons showed reliable outward photocurrents (Fig. 1a) in response to 470 nm full field light pulses. The photocurrent after 1 s of continuous illumination (stationary photocurrent) of GtACR2 expressing neurons clamped to -35 mV was significantly higher than that of the engineered ACRs (eACRs) iC++ and iChloC (628.5 ± 61.8 pA, 330.2 ± 37.9 pA, and 136.3 ± 21.4 pA, respectively; Fig. 1b). Given the poor membrane targeting and intracellular accumulation of GtACR2 (Fig. 1c), the high single-channel conductance of GtACR2 ^16^ is likely the cause for the high photocurrents observed in the whole-cell recordings. While the native GtACR2 seems to outperform the previously-described eACRs, the above findings suggest that improved membrane targeting of GtACR2 would greatly facilitate silencing of neuronal activity. Importantly, this should allow efficient silencing by using significantly lower light power, enabling optogenetic control of a larger brain volume.

**Figure 1.**
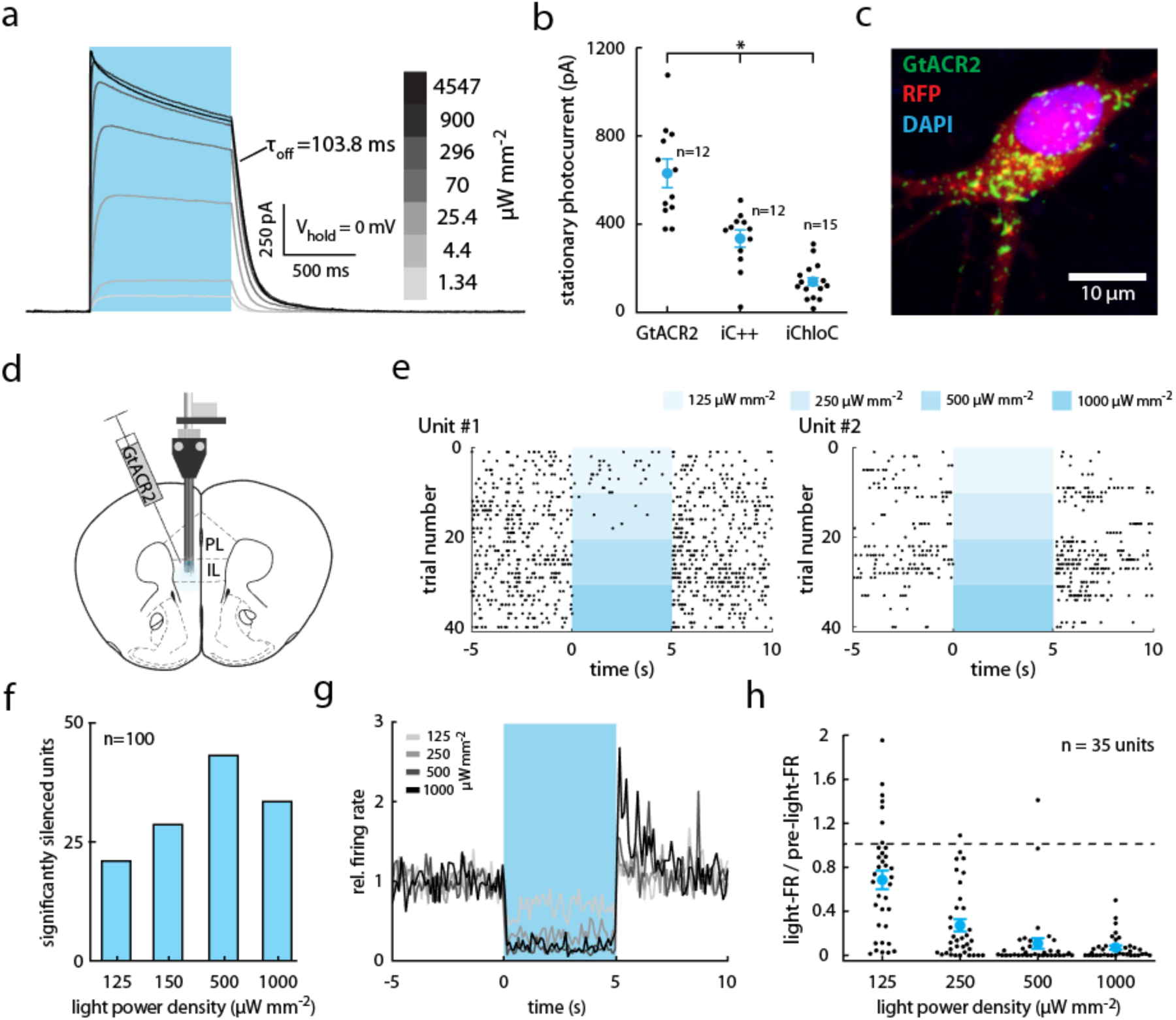
GtACR2 efficiently silences neuronal activity *in vivo*. **(a-c)** Characterization of ACRs in cultured rat hippocampal neurons. **(a)** Sample whole-cell voltage-clamp photocurrent recording of a GtACR2-expressing cell illuminated (470 nm) with increasing light power density. **(b)** Comparison of stationary photocurrents of blue light-sensitive ACRs (V_hold_ = -35 mV, current after 1 s of continuous illumination). Neurons expressing GtACR2 (n = 12) showed the highest photocurrents compared with neurons expressing iC++ (n = 12) and iChloC (n = 15). F(2,36) = 36.92, P = 1.9 × 10^-9^ **(c)** Representative image of GtACR2 localization. Green: GtACR2, red: cytoplasmic RFP, blue: nucleus. **(d-h)** *In vivo* quantification of GtACR2 mediated neuronal silencing efficiency. **(d)** Schematic of experimental paradigm. Extracellular recordings from the mPFC were performed with a movable optrode following injection of AAV2/1 encoding GtACR2 into the mPFC. **(e)** Two representative raster plots of units significantly reducing their firing rate during 5 s of illumination with blue light (460 nm). Each light power was tested 10 times. While the unit depicted on the left shows a graded response to increasing light powers, the unit depicted on the right is completely inhibited even at the lowest tested light power. **(f)** Number of units that significantly reduced their firing rate compared to 5 s pre-light period, dependent on the tested light powers. **(g)** Normalized firing rate (FR / pre-light-FR, 100 ms bins) of all significantly silenced units. **(h)** Quantification of g. In b and h, results are presented as means (± SEM).

To verify this prediction, we characterized the efficiency of GtACR2-mediated inhibition in awake, behaving mice with extracellular recordings (Fig. 1d-g). To quantify the efficiency of silencing in a large cortical volume, we recorded from mice expressing GtACR2 in pyramidal neurons in the medial prefrontal cortex (mPFC). Mice were implanted with movable fiberoptic-coupled microwire arrays in which electrodes were placed 500 μm below the optical fiber tip (Fig. 1d). Using analytical modeling of light scattering and absorption in brain tissue ^23^ we estimated the light power density at the position of the extracellular recording site to be 0.11% of the light power density exiting the optical fiber (Supplementary Fig. 1). Single-unit recordings showed a significant reduction in neuronal firing rates in response to a 5 s 460 nm light pulse (Fig. 1e-h). These recordings showed that out of 100 single units recorded (*n* = 2 mice, 12 recording sites), 43% and 35% significantly reduced their firing rate at the second highest and highest tested light power densities (0.5 and 1 mW mm^-2^, respectively; Fig. 1f). Half of the silenced units showed a significant firing rate reduction already at the lowest tested light power density (125 μW mm^-2^), with some units completely silenced (Fig. 1e, h). At the highest light intensity, silenced units showed pronounced rebound activity upon light pulse termination (Fig. 1g). Notably, all light power densities used were well below the necessary light powers for *in vivo* optogenetic inhibition using microbial ion-pumps ^8,6^, indicating that GtACRs can serve as potent inhibitory optogenetic tools for somatic silencing in mammalian neurons.

### Activation of ACRs in the axonal compartment induces antidromic action potentials

We have previously shown that GtACR1 can induce vesicle release from thalamocortical projection neurons upon illumination of their axonal terminals in the acute brain slice ^13^. Malyshev and colleagues 24 demonstrated a similar effect in GtACR2-expressing cortical pyramidal neurons in acute brain slices. To verify that this excitatory action of GtACRs is not an artifact of the acute brain slice preparation, we evaluated the excitatory effect of ACRs in two separate preparations: in cultured hippocampal neurons and in awake, freely moving mice. Whole-cell patch-clamp recordings in current-clamp mode from GtACR2-expressing cultured neurons revealed that during a 100 ms long illumination pulse, GtACR2 reliably inhibited action potential (AP) generation (Fig. 2a). However, this was often associated with what appeared to be an attenuated AP shortly after light onset (‘escaped AP’, Fig. 2a, inset). When recorded in voltage-clamp mode, escaped APs were measured in 6 out of 12 tested GtACR2-expressing neurons in response to 1 ms light pulses at saturating light power (4.5 mW mm^-2^). These escaped spikes occurred even when the recorded photocurrent was an outward current, expected to hyperpolarize the somatic membrane (Fig. 2b, upper left). We then asked whether these light-evoked antidromic APs are specific to naturally-occurring chloride-channels, or a general feature of light-evoked chloride conductance in the axon. We therefore tested two engineered anion-conducting channelrhodopsins under the same experimental conditions. APs were evoked in response to 1 ms and 1 s long light pulses in 2 / 13 and 7 / 12 iC++ expressing neurons, respectively, showing that this effect is not specific to GtACRs (Fig. 2b, middle). No APs were evoked in iChloC expressing cells (n = 22 and n = 15, for 1 ms and 1 s long light pulses, respectively; Fig. 2b right), likely because of the overall smaller photocurrents we observed in cells expressing this construct (Fig. 1b). To further test the hypothesis that ACR activation might be depolarizing in some neuronal compartments, we applied spatially-restricted laser pulses to the soma or neurites of cultured hippocampal neurons during whole-cell patch-clamp recordings. Light pulses directed at the soma using a galvanometric mirror system (see Online Methods) induced small hyperpolarizing or depolarizing photocurrents, while light pulses directed to neurites of the same cell evoked antidromic spikes (Fig. 2c).

**Figure 2.**
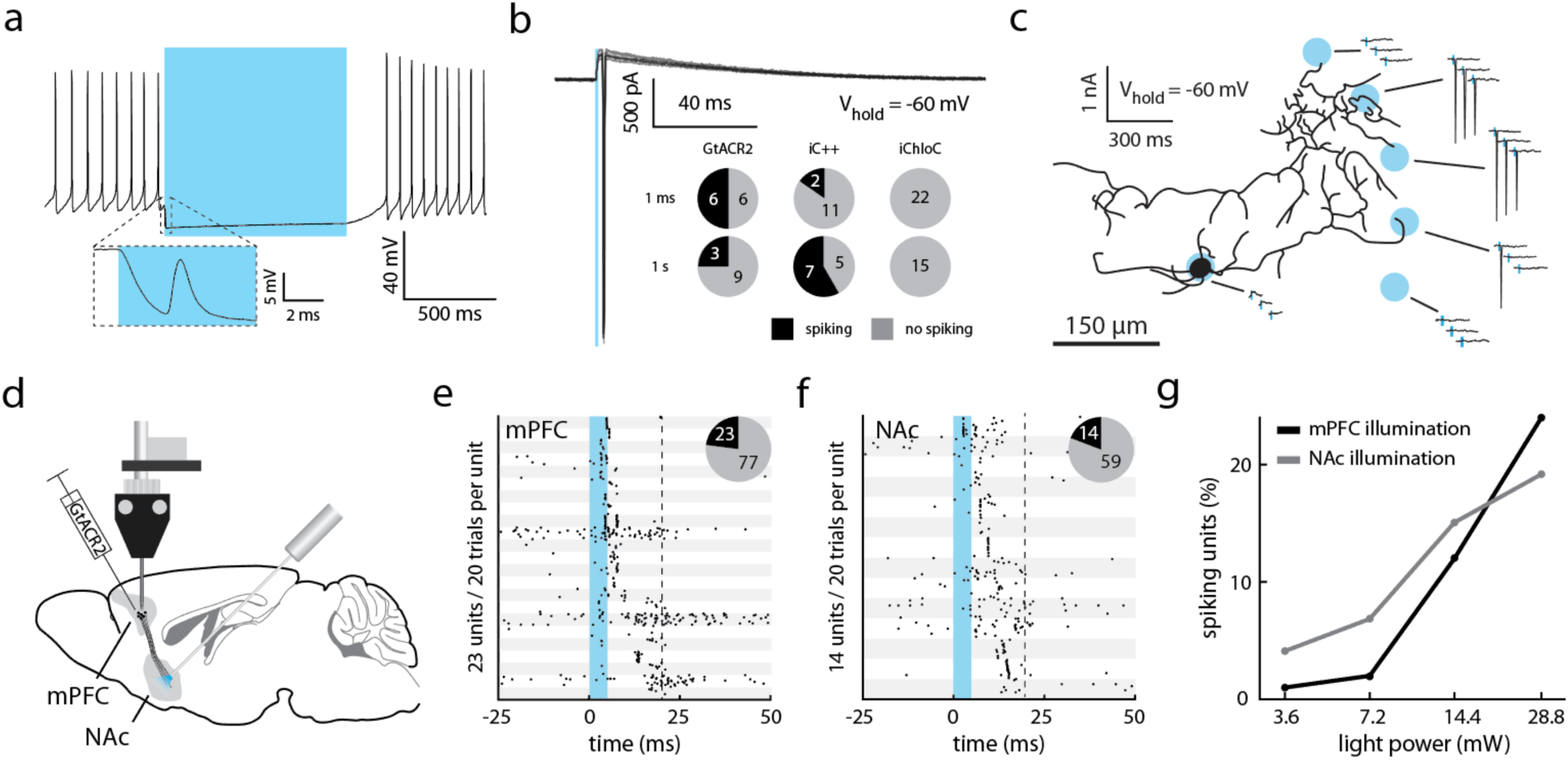
Activation of GtACR2 in the axonal compartment induces action potentials *in vitro* and *in vivo*. **(a-b)** Characterization of light-evoked spiking in ACR-expressing cultured hippocampal neurons (470 nm at 4.5 mW mm^-2^). **(a)** Representative whole-cell current-clamp recording of a GtACR2-expressing cell silenced by light application. Inset: strongly attenuated spike occurring shortly after light onset. **(b)** Representative whole-cell voltage-clamp recording of escaped action potentials in response to 1 ms light pulses. Pie charts depict the number of neurons with induced spikes for the three tested light-gated chloride channels: GtACR2, iC++ and iChloC. (**c**) Illumination of distal neurites induces spiking in cultured neurons. Schematic depicting the outline of a GtACR2-expressing neuron overlaid with the locations of laser illumination spots. Shown are whole-cell voltage-clamp responses to spatially-restricted illumination at the indicated locations. **(d-g)** *In vivo* extracellular recording following GtACR2 expression in the mPFC. **(d)** Schematic of the implantation allowing for illumination of the NAc, a downstream target of the mPFC, while recording in the mPFC. (**e**) Single units recorded in the mPFC, showing rapid light-evoked responses during a 20 ms time-window starting with a 5 ms light pulse (1 mW mm^-2^, corresponding to 28.8 mW at the fiber tip). Units are arranged from top to bottom according to their mean first spike latency across 20 trials. Pie charts depict the number of neurons with significantly increased spike rates. **(f)** mPFC units showing significantly increased firing rates in response to illumination of the NAc. Units are sorted by mean spike latency. Light power at the fiber tip: 28.8 mW. Pie chart as in (e). **(g)** Percent of units with increased firing rate in response to 5 ms light pulses of increasing light power.

These recordings were performed in cultured neurons after at least 14 days *in vitro*, a stage at which intracellular chloride concentrations should reach the adult state ^25^ as the expression of the neuronal potassium chloride co-transporter KCC2 ^26^ is fully up-regulated ^27^. Nevertheless, chloride homeostasis of neuronal cultures may differ due to reduced concentrations of KCC2 regulators such as insulin ^28^ in the culture medium. We therefore tested whether activation of GtACR2 could lead to axonal excitation *in vivo.* We recorded from GtACR2-expressing mice (Fig. 1d) using movable fiberoptic-coupled microwire drives at the site of AAV injection (Fig. 1d-h). In the same animals, we implanted a second optical fiber terminating at the nucleus accumbens (NAc), a prominent projection target of the mPFC ^29^ (Fig. 2d). In response to brief light pulses to the mPFC (5 ms pulse width, 460 nm at 1 mW mm^-2^, which corresponds to 28.8 mW at the fiber tip), APs were evoked (Fig. 2e,g) in the same AAV-expressing region that showed significant silencing during 5 s light pulses (Fig. 1d-h). Moreover, similar 5 ms light pulses delivered to the NAc led to the induction of short-latency APs in the mPFC (Fig. 2f-g), presumably due to axonal excitation and antidromic propagation.

In summary, light stimulation of GtACR2-expressing axons led to APs in hippocampal neurons *in vitro,* in thalamocortical projection neurons in acute brain slices ^13^ and in striatum-projecting cortical neurons in awake, behaving mice. We therefore concluded that chloride-mediated axonal depolarization is a general phenomenon that could confound the analysis of optogenetic silencing experiments.

### Overexpression of KCC2 reduces GtACR2-mediated antidromic action potentials

Based on these findings, we reasoned that if GtACR-mediated antidromic spiking is indeed due to a positively shifted chloride reversal potential in the axon, decreasing the axonal chloride concentration should reduce the probability of antidromic spike generation. The chloride extruder KCC2^26^ is up-regulated in neurons during development, leading to high endogenous KCC2 protein levels in somatic and dendritic membranes ^30^, but does not localize to axons^31,32,33^. We first tested whether overexpression of KCC2 leads to its localization to the axonal compartment in cultured hippocampal neurons. We co-transfected neurons with expression vectors encoding the green fluorescent protein mNeonGreen (GFP) and KCC2, or with GFP alone as control. We then labeled the neurons with antibodies against the dendrite-specific microtubule-associated protein-2 (MAP2) and KCC2. Overexpression of KCC2 led to strong KCC2 immunoreactivity in dendrites and somata of transfected neurons (Fig. 3a), compared to endogenous expression levels (Fig. 3a, white arrow). To quantify axonal KCC2 levels, axons were detected as neurites that are GFP-positive and MAP2-negative (Fig. 3a, zoom in). While mean axonal KCC2 intensity was not significantly different between young (7 days *in vitro*) and mature (16 days *in vitro*) hippocampal cultures, KCC2 overexpression led to a 6.6 ± 1.1 fold higher axonal KCC2 signal (Fig. 3b, ctrl young vs. ctrl mature: P = 0.20; ctrl young vs. KCC2: P = 5*10^-7^; ctrl mature vs. KCC2: P = 9.8*10^-3^). KCC2 expression did not significantly shift the chloride reversal potential measured in the soma (Fig. 3c) or the action potential initiation threshold (rheobase, Fig. 3d), but indeed led to a significant reduction of GtACR2-evoked antidromic spiking (1 ms light pulses width, 470 nm at 4.5 mW mm^-2^; Fig. 3e). This result provides further support for the notion that the chloride reversal potential in the axon is depolarizing under physiological conditions.

**Figure 3.**
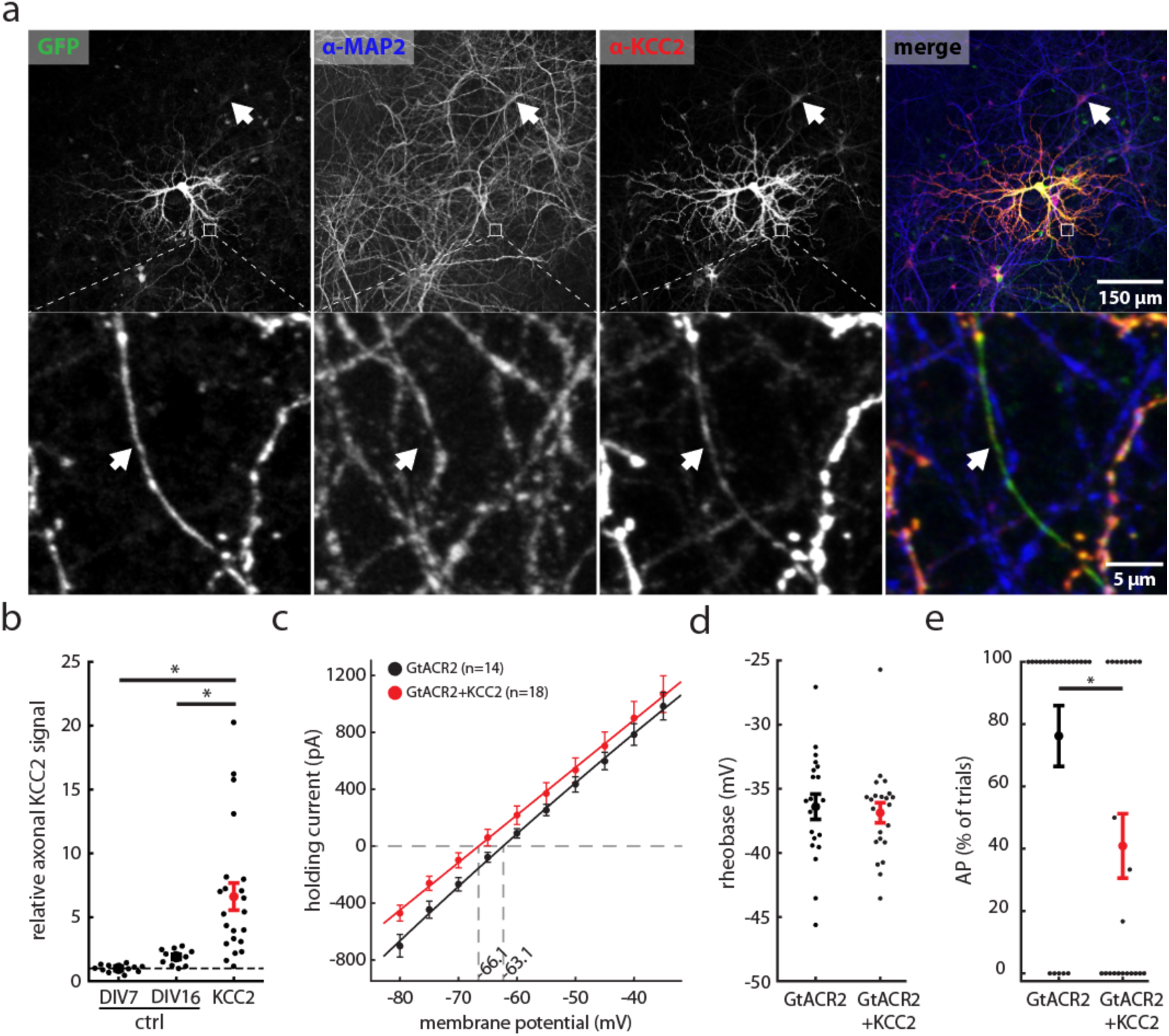
Overexpression of KCC2 reduces antidromic spiking in cultured hippocampal neurons. **(a)** Images from an KCC2 overexpression experiment. Endogenous KCC2 is expressed in the somatic compartment, while overexpression of KCC2 led to increased expression in axonal projections. Cultured hippocampal neurons were sparsely transfected either with GFP alone (ctrl) or GFP and KCC2. Neurons were then fixed and stained for MAP2 and KCC2. Top images show a representative region of interest with one overexpressing cell in the center. The arrow indicates a neuronal cell body expressing endogenous KCC2 levels at 16 days *in vitro* (DIV). Bottom images depict MAP2-expressing dendrites and a single MAP2-negative axon (arrow), which is positive for overexpressed KCC2 based on its anti-KCC2 fluorescence. **(b)** Quantification of axonal KCC2 immunofluorescence for immature (DIV7) and mature control neurons (DIV16) and neurons overexpressing KCC2, normalized to the average axonal KCC2 signal in immature neurons (DIV7). Axonal KCC2 fluorescence is significantly higher in KCC2 overexpressing cultures. (Kruskal–Wallis H test, H(2,44) = 29.26, P < 10^-4^; ctrl: n_DIV7_ = 11, n_DIV16_ = 10, KCC2: n = 23) **(c-e)** Physiological properties and light-evoked spiking in cultured hippocampal neurons expressing either only GtACR2, or co-expressing KCC2. **(c)** Effect of KCC2 overexpression on the IV-curve. The reversal potential did not differ significantly (Students t-test = 1.5, GtACR2: n = 14, GtACR2+KCC2: n = 18, P = 0.15 two-tailed). **(d)** Comparison of the minimal current injection to induce an action potential (rheobase). KCC2 overexpressing neurons did not differ from GtACR2 only expressing neurons (Students t-test = 0.5, GtACR2: n = 21, GtACR2+KCC2: n = 22, P = 0.7 two-tailed). **(e)** KCC2 overexpression significantly reduced the likelihood of GtACR2 mediated action potential generation. (Mann–Whitney U = 146.5 n_GtACR2_ = 21 n_GtACR2+KCC2_ = 22, P = 4 * 10^-2^).All results are presented as means (± SEM).

### Soma-targeting of GtACR2 increases somatic photocurrents and reduces axonal excitation

To overcome the two main caveats of GtACRs in respect to their utility as optogenetic inhibitory tools, namely their poor membrane targeting and triggering of antidromic spikes in the axonal compartment, we designed several new GtACR2 variants with altered membrane targeting sequences (Fig. 4a). Addition of the ER export and trafficking signals from the mammalian inward rectifying potassium-channel Kir2.1 was previously shown to reduce intracellular aggregation of the chloride-pump NpHR ^34,35^. Indeed, fusion of these sequences to GtACR2 (eGtACR2) led to reduced intracellular accumulation *in vivo* (Fig. 4c, 2^nd^ column). To reduce antidromic spike generation, GtACR2 should be removed from the axonal compartment. To achieve soma-specific localization of GtACR2, we replaced the ER export signal with the soma-targeting motif of the soma and proximal dendrite localized voltage-gated potassium-channel Kv2.1 ^36,37^, which was previously shown to enhance soma-localized expression of channelrhodopsin-2 ^38,22^. Hypothesizing that destabilizing the protein by adding a protein degradation-promoting proline (P), glutamic acid (E), serine (S), and threonine (T) rich sequence (PEST ^39^) could limit the effective lifetime of membrane-resident channels diffusing along the axon, thereby potentially further restricting GtACR2 protein levels outside the somatic compartment. To test these new viral constructs, we infused AAVs encoding them unilaterally into the mPFC of mice, leading to strong fluorescence at the injection site as well as sparse labeling along the injection needle track (Fig. 4b). Soma-targeted GtACR2 (stGtACR2; Fig. 4c, 3^rd^ column) as well as the destabilized stGtACR2-PEST (Fig. 4c, 4^th^ column) showed improved membrane targeting, strong soma-associated fluorescence and reduced neurite fluorescence (Fig. 4d). Functional characterization of the soma-targeted constructs by whole-cell patch-clamp recordings in the acute brain slice showed a 2.6-fold increase in stationary photocurrents compared to untargeted GtACR2, leading to average photocurrents of more than 2 nA when cells were clamped to -35 mV (Fig. 4e). Improved membrane targeting alone strongly increased the antidromic spike generation probability (Fig. 5a; eGtACR2), while soma-targeting not only increased photocurrents (Fig. 4e) but also decreased the probability of inducing antidromic spikes in cultured hippocampal neurons (Fig. 5a). Destabilizing stGtACR2 using the PEST sequence led to a less pronounced reduction in antidromic spike generation compared to stGtACR2 (Fig. 5a). Photocurrents were quantified in the same neurons to verify that the reduced probability of antidromic spike generation is not due to differences in peak photocurrents of the different constructs in cultured neurons (Fig. 5b). In contrast to the stationary photocurrents in acute brain slice experiments (Fig. 4e) peak photocurrents in cultured neurons did not differ significantly between constructs, pointing to a lower membrane targeting efficiency in cultured neurons or an influence of the shorter virus incubation time. Nevertheless, it follows that the dramatic reduction in antidromic spiking for stGtACR2 is not due to lower photocurrents.

**Figure 4.**
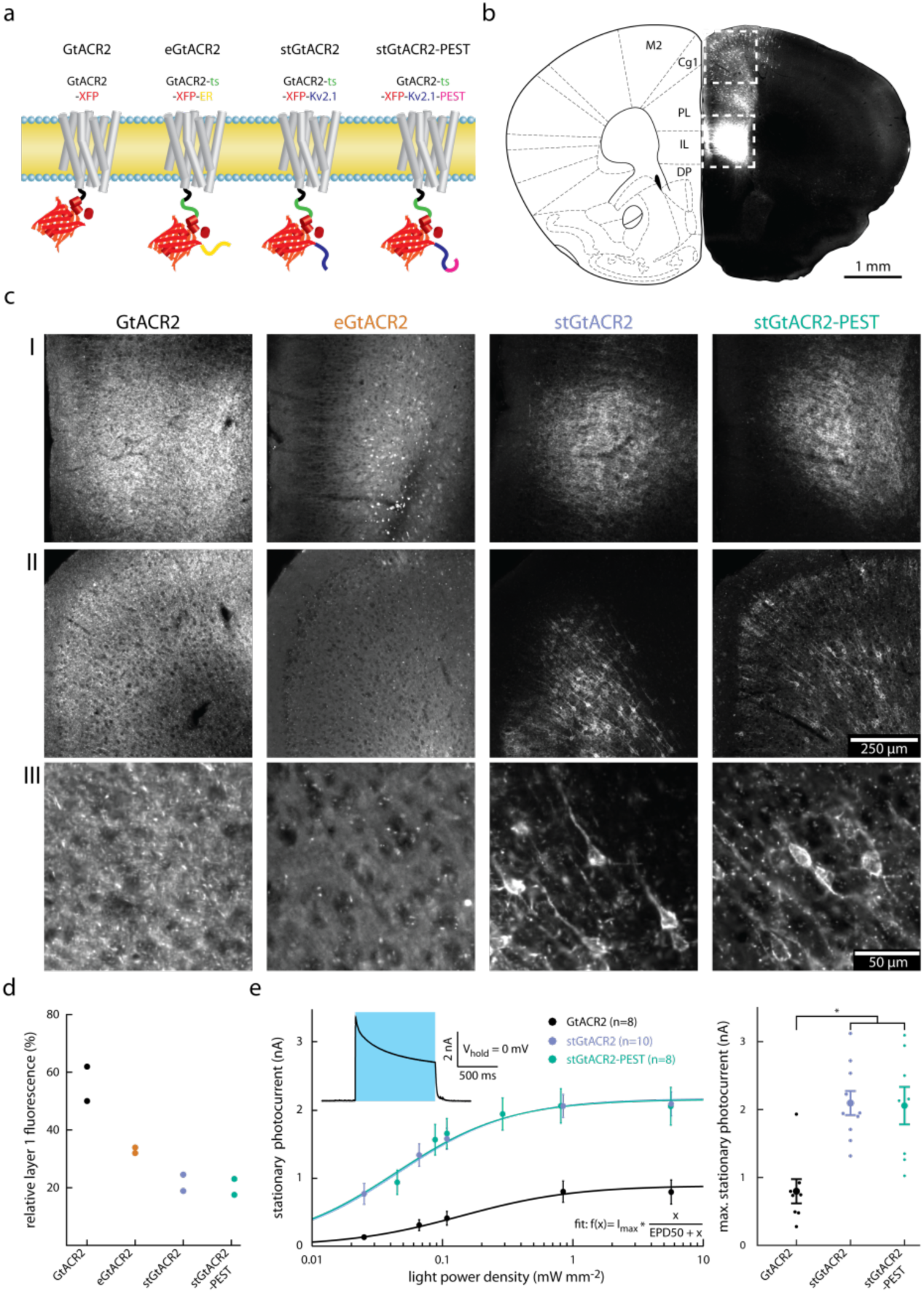
Targeting GtACR2 to the neuronal soma leads to enhanced photocurrent amplitude. **(a)** Schematic of different targeting approaches. **(b)** Image showing the fluorescence resulting from AAV2/1 mediated cytosolic fluorophore expression in the mPFC. Transduction is most dense at the injection site (indicated by the lower dashed box) and sparse along the injection needle track (upper dashed box). **(c)** Higher magnification images of the areas indicated in b. c-I: Zoom in on the injection site. c-II: Zoom in on the more dorsal region of sparse expression. c-III: Higher magnification of c-II. stGTACR2 and stGtACR2-PEST show enrichment at the soma. **(d)** Quantification of soma restriction by normalizing mPFC layer 1 fluorescence by the mean fluorescence measured at the injection center. Targeting reduces relative layer 1 fluorescence (n = 2 per group). **(e)** Light power density dependence of stationary photocurrent in whole-cell patch-clamp recordings of neurons in acute brain slices. Inset: Representative whole-cell voltage-clamp recording. The stationary photocurrent was defined as the photocurrent at the end of a 1 s light pulse. The fit was performed per cell with the effective light power density for 50% photocurrent (EPD50) as free parameter. stGtACR2 and stGtACR2-PEST have a significantly higher maximal stationary photocurrent than GtACR2 (F(2,23) = 11.84, P = 2.9 × 10^-4^; GtACR2: n=8, stGtACR2: n = 10, stGtACR2-PEST: n = 8). Results are presented as means (± SEM).

**Figure 5.**
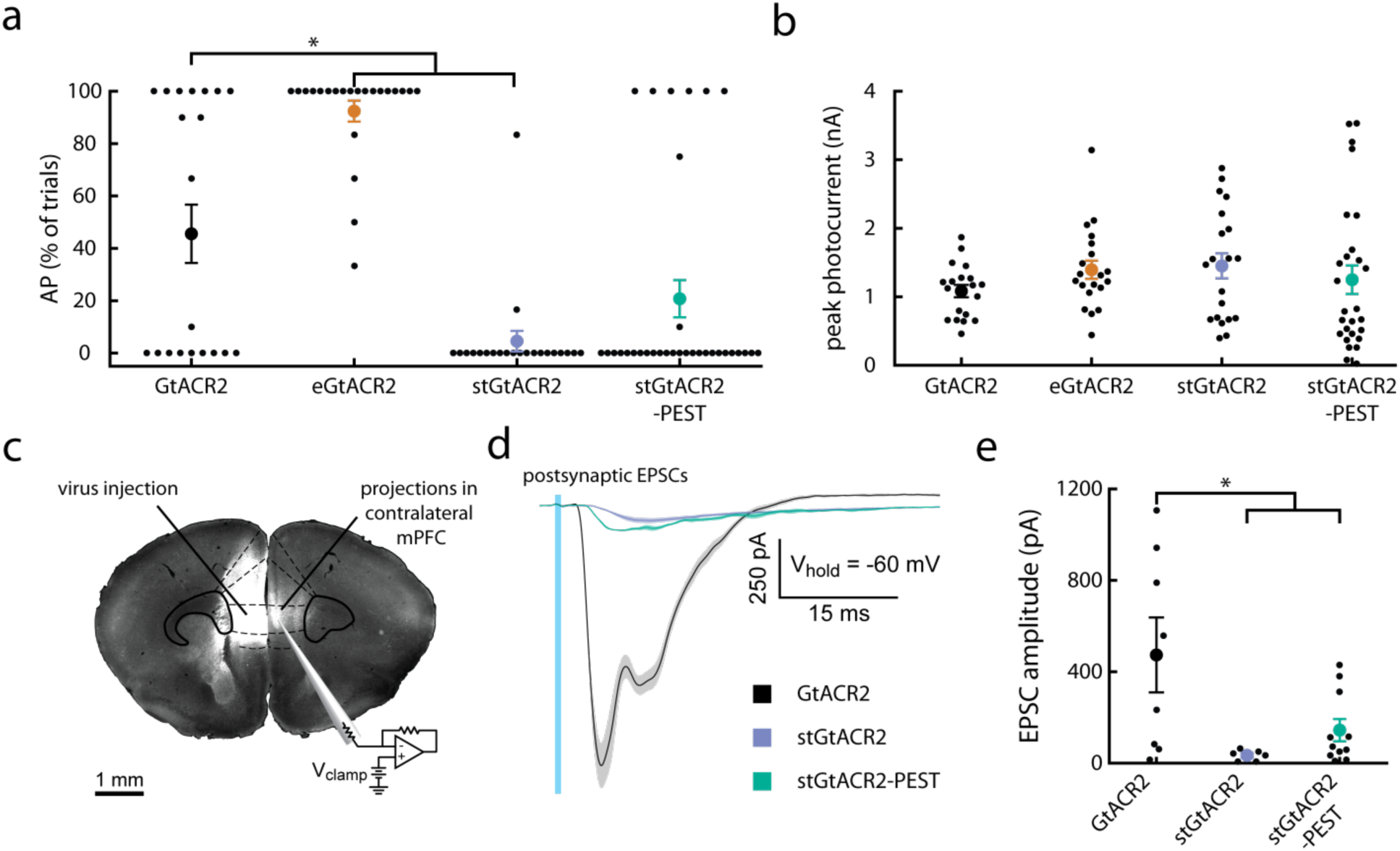
Targeting GtACR2 to the somatodendritic compartment attenuates axonal excitation. **(a)** Comparison of the incidence of antidromic spikes triggered by 1 ms light pulses in virally transduced cultured hippocampal neurons. eGtACR2 has a significantly increased AP incidence, while stGtACR2 decreases the occurrence of APs. (Kruskal–Wallis H test, H(3,97) = 47, P < 10^-4^; n GtACR2 = 20, n eGtACR2 = 22, n stGtACR2 = 22, n stGtACR2-PEST = 33) **(b)** Peak photocurrents did not differ significantly between the constructs, showing that reduced AP incidence in stGtACR2 transduced neurons does not stem from smaller photocurrents. (F(3,82) = 0.83, P = 0.48). **(c)** Schematic of the experimental setup to characterize GtACR2 triggered axonal neurotransmitter release in acute brain slices. Virus encoding a cytosolic fluorophore was co-injected with the GtACR2 variants to allow for visualization of the axon terminals of transduced cortico-cortical projection neurons. Contralateral neurons in areas with high fluorescence intensity were recorded. **(d)** Representative traces of excitatory post-synaptic currents in response to 1 ms light pulses (470 nm, at 4.5 mW mm^-2^). **(e)** Quantification of the light evoked post-synaptic current amplitude. Soma targeting led to significant reduction in light evoked EPSCs amplitudes (F(2,22) = 5.54, P = 1.13*10^-2^). All results are presented as means (± SEM).

We next asked whether stGtACR2 and stGtACR2-PEST would show improved performance through reduced light-evoked synaptic release from long-range projecting axons. We injected AAVs encoding the GtACR2 variants together with a second AAV encoding a cell-filling fluorophore unilaterally to the mPFC, to allow for visualization of the axons of cortico-cortical projection neurons in the contralateral hemisphere (Fig. 5c). During acute brain slice preparation from these mice, the corpus callosum was severed, separating the somata of the transduced cortico-cortical projecting neurons from their axon terminals in the contralateral mPFC. Conducting whole-cell patch-clamp recordings from postsynaptic neurons in areas with fluorescently-labeled axons contralateral to the injection site therefore allowed us to characterize the isolated effect of GtACR2 activation on the axonal compartment. Blue light pulses led to reliably evoked EPSCs in slices expressing the non-targeted GtACR2 (Fig. 5d). In contrast, the EPSC amplitude was dramatically reduced in slices expressing the soma-restricted stGtACR2 (Fig. 5d,e; 473.4 ± 153.4 pA vs. 34.1 ± 9.5 pA for GtACR2 and stGtACR2, respectively). The increased somatic photocurrents of stGtACR2, together with the near-elimination of antidromic spiking and neurotransmitter release, make it a highly efficient tool for optogenetic inhibition.

### stGtACR2 mediated BLA inhibition prevents fear extinction learning

To verify the utility of stGTACR2-mediated optogenetic inhibition in awake, behaving animals, we chose to use this tool for suppressing basolateral-amygdala (BLA) activity during extinction of auditory-cued fear conditioning, a well-established form of associative learning ^40^. The BLA plays a central role in the acquisition as well as extinction of the conditioned freezing response. Based on previous work ^41^, we hypothesized that temporally-precise inhibition of the BLA during the delivery of conditioned stimuli in extinction training would suppress the formation of extinction memory. We bilaterally injected mice with AAV encoding stGtACR2 or a fluorophore-only control vector into the BLA and implanted 200 μm-diameter optical fibers above the injection sites (Fig. 6a; Supplementary Fig. 2). Following 3 weeks of recovery, mice underwent fear conditioning in context A (Fig. 6b). Both groups (stGtACR2, n = 8; control, n = 8) showed increased freezing during acquisition (ctrl: from 3.8±1.9% to 39.5±8.1%; stGtACR2: from 3.5±1.5% to 29.6±3.5%, Scheirer Ray Hare test H = 52.91, P = 3.5*10^-10^) with no significant difference between groups (ctrl vs. stGtACR2: Scheirer Ray Hare test H = 0.11, P = 0.74), suggesting that BLA activity is not altered merely by expression of stGtACR2 (Fig. 6c). To test for fear recall and extinction, mice underwent extinction training two days later in a different context from that in which they were fear conditioned (context B). The extinction protocol consisted of twenty 30 s tone presentations that were paired with blue light delivery (447 nm; 5 mW from each fiber tip). Mice were then tested in an extinction retrieval test the following day in which they were subjected to twenty CS presentations, but no light was delivered (Fig. 6b, right). During this test, stGtACR2 mice showed higher freezing rates during CS presentation (ctrl vs. stGtACR2: Scheirer Ray Hare test H = 4.30, P = 3.8*10^-2^), but freezing levels were indistinguishable from control mice during the inter-tone intervals (ctrl vs. stGtACR2: Scheirer Ray Hare test H = 3.6*10^-2^, P = 0.85), indicating that fear extinction was prevented by stGtACR2 mediated BLA inhibition during CS presentation. Sierra-Mercado and colleagues ^41^, previously showed that inhibition of the BLA by muscimol injection prior to fear extinction interfered with extinction learning. Our results extend these findings, demonstrating that temporally-precise inhibition of BLA activity only during CS presentation using stGtACR2 can interfere with extinction learning. In summary, our experiments indicate that stGtACR2 is a powerful inhibitory optogenetic tool, allowing temporally precise silencing of neuronal populations *in vivo.*

**Figure 6.**
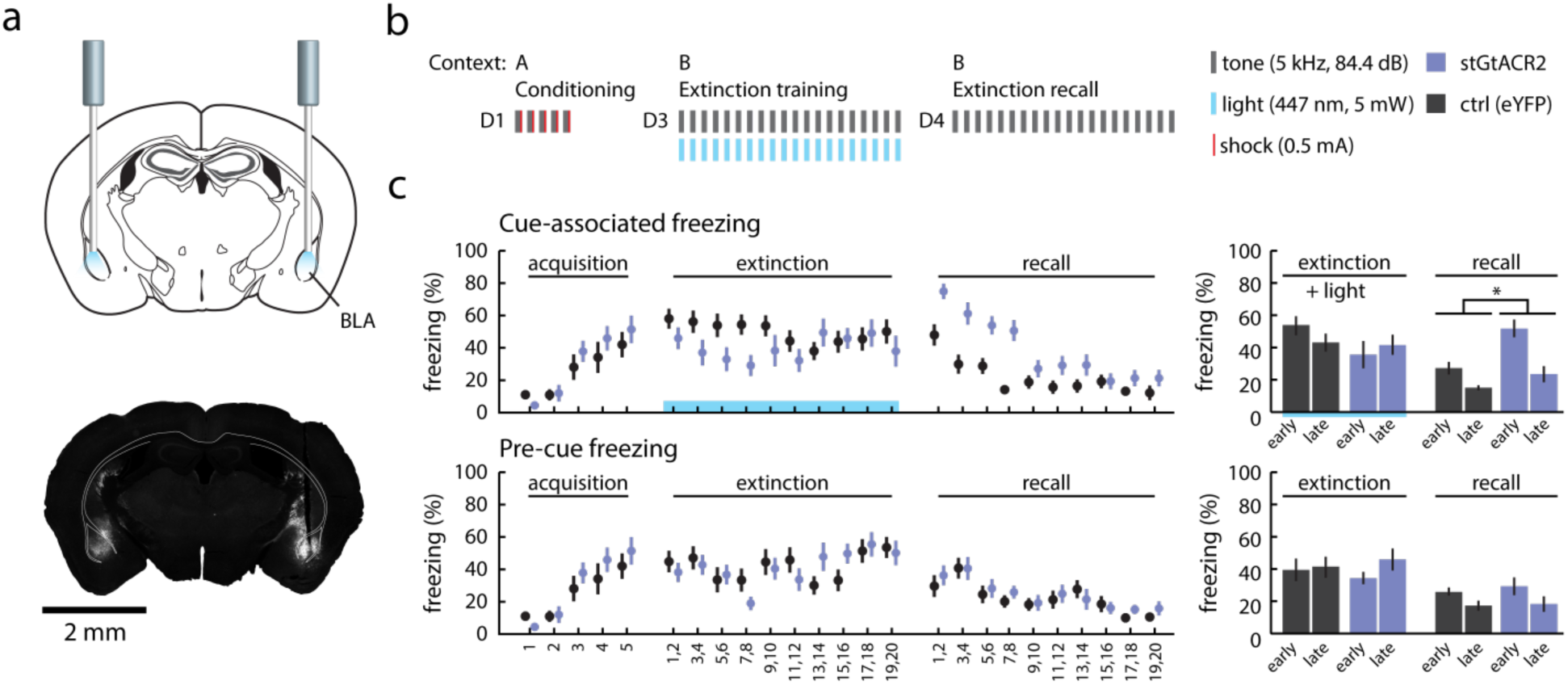
Silencing of cue-associated BLA activity using stGtACR2 suppresses extinction of cued freezing. **(a)** Mice were bilaterally injected with eYFP or stGTACR2-encoding virus and implanted with optic fibers targeting the BLA. Bottom: representative image of stGtACR2 expression in the BLA. **(b)** stGtACR2 and control (eYFP) mice were subjected to auditory fear conditioning (day 1, conditioning), extinction training (day 3, early and late extinction) and extinction recall (day 4, early and late recall). During auditory fear conditioning mice were submitted to five tone (CS)-shock (US) presentations in context A. On day 3 twenty 30 s tone (CS) presentations were paired with light (447 nm at 5 mW exiting the fiber tip) in context B. On day 4 extinction recall was tested by twenty 30 s tone presentations in context B. **(c)** Percentage of freezing during presentation of the CS (top row) and the 30 s prior to CS (bottom row). The right column depicts the mean percentage of freezing during the early and late phases of the trials for the two groups. Mean freezing levels for ctrl and stGtACR2 mice, during the early recall phase, were 28 ± 4 % and 54 ± 6 %, respectively; the distributions in the two groups differed significantly (Scheirer Ray Hare test H = 4.30, P = 3.8*10^-2^). All results are presented as means (± SEM).

## Discussion

We took a membrane targeting approach to allow the utilization of the high-conductance *Guillardia theta* anion-conducting channelrhodopsins ^16^ as an optogenetic tool in mammalian neurons. While these naturally-occurring channelrhodopsins showed great promise due to their highly efficient photocurrents and light sensitivity ^16^, and have proven effective in silencing drosophila and zebrafish neurons ^19,20,21^, they have seen little use in mammalian neuroscience applications. This was mainly due to poor membrane targeting and to complex effects on axonal excitability ^13,24^. Our findings indicated that even in its non-targeted form, GtACR2 can efficiently silence neurons in the medial prefrontal cortex of behaving mice. These results were consistent with the high photocurrent amplitudes recorded in neurons expressing GtACR2, compared with cells expressing the engineered ACRs iC++ and iChloC ^17,15^. Given the high single channel conductance, favorable photon-ion stoichiometry, and high light sensitivity, the light power density for neuronal inhibition with GtACR2 is at least one order of magnitude lower than that of other silencing opsins ^8,42^. However, despite its apparent high efficacy, a significant portion of the protein seemed to reside in intracellular compartments, where it cannot contribute to functional photocurrents. Furthermore, activation of GtACR2 in our recordings was also associated with antidromic spiking at light onset when illuminating both the proximal and distal axons. We have previously observed GtACR2-mediated triggering of synaptic release in thalamocortical axons^13^, consistent with recent reports of antidromic spiking in layer 2/3 pyramidal neurons^24^ in the slice preparation. In this study, we observed GtACR2-mediated antidromic spiking in cultured hippocampal neurons, in cortico-cortical neurons in the acute slice and in cortico-striatal axons of behaving mice. Our findings indicate that axonal excitation by a chloride conductance is a general phenomenon, and could reflect a depolarized reversal potential for chloride in the axonal compartment. While such effects have been previously reported, for example in hippocampal mossy fibers^43,44^, cerebellar^45^ and brain-stem axons^46,47^, systematic evaluation of the phenomenon has been previously restricted to axons that naturally express GABA-A receptors.

To determine whether elevated chloride concentration in the axon could indeed lead to GtACR2-mediated axonal excitation, we co-expressed the KCC2 transporter with GtACR2 in cultured neurons. The endogenous KCC2 transporter, which is expressed in mature neurons and is known to be responsible for extruding chloride from the somatodendritic compartment^48^, is known to be absent from the axon ^31,32,33^, potentially permitting a higher chloride concentration in this compartment. Our finding that overexpression of KCC2 resulted in a significant decrease in light-induced antidromic spiking indicates that ACR-mediated antidromic spiking could indeed be the result of a smaller chloride gradient in the axon, even in adult neurons. While this antidromic spiking phenotype would probably not interfere with long-term inhibition experiments (minutes and upward), it might be a confounding factor when temporally-precise (millisecond-scale) inhibition is required. Future work could combine GtACR2 stimulation with red-shifted chloride indicators to directly examine changes in chloride levels in the axonal compartment during ACR-mediated chloride conductance.

Most importantly, our study demonstrates that the soma-targeted variants of GtACR2 show improved membrane expression in the somatodendritic compartment, and offer superior anion photocurrents for high-efficiency optogenetic silencing of neurons in the mammalian brain. Current optogenetic experiments often involve a sparsely labeled population of neurons that are distributed across a large brain tissue volume^49,50^. Efficient silencing of such widely-distributed neuronal populations require continuous activity of the inhibitory optogenetic tool^51^, placing considerable constraints related to tissue heating and photodamage^52,9^. Our data indicate that stGtACR2 can provide an effective means of performing such challenging experiments, due to its intrinsically high conductance, which increases its effective light sensitivity in expressing neurons. With increasing distance from the fiber tip, the wavelength dependent transmittance becomes increasingly relevant. For instance, multiplying the action spectra of GtACR1 and GtACR2 ^16^ with the analytically modeled light transmittance curve ^23^ for brain tissue (Supplementary Fig. 1) revealed that excitation of GtACR2 with 480 nm or of GtACR1 with 510 nm would provide optimal light-mediated silencing at 500 μm distance from the optic fiber tip. In experiments that require optogenetic manipulation of functionally- but not anatomically-segregated neuronal populations, stGtACR2 might be combined with red-shifted tools such as C1V1 ^53^, Chrimson ^54^ or ReaChR ^55^. Red-shifted calcium sensors ^56,57^ could also be used in combination with stGtACR2 due to its minimal responsivity at wavelengths above 560 nm. Notably, both stGtACR1 and stGtACR2 are also highly advantageous for multiphoton single-cell silencing experiments ^58,59^ owing to their somatic restriction ^22^ and high-amplitude photocurrents.

In summary, we have demonstrated that membrane targeting and somatodendritic restriction of the naturally-occurring anion-conducting GtACR2 address two independent constraints of this channelrhodopsin, greatly improving photocurrents and minimizing axonally-generated antidromic action potentials. We were able to achieve high-efficiency neuronal silencing with the optimized stGtACR2 and demonstrated its efficacy for temporally-precise inhibition of amygdala activity during extinction learning.

## Acknowledgments

We thank M. Segal (Weizmann Institute of Science) for reagents. We thank R. Zwang for help with cloning. We thank the Yizhar laboratory members for comments on the manuscript. We acknowledge support (to O.Y.) from the Human Frontier Science Program, a European Research Council starting grant (ERC-2013-StG OptoNeuromod 337637), the Adelis Foundation, the Lord Sieff of Brimpton Memorial Fund and the Candice Appleton Family Trust. O.Y. is supported by the Gertrude and Philip Nollman Career Development Chair.

## Author Contributions

M.M. and O.Y. designed the study with input from I.L.; M.M. designed the constructs, performed *in vitro* electrophysiology and imaging experiments. K.CKM. performed *in vitro* neuronal recordings. L.G. performed and analyzed *in vivo* electrophysiology recordings. P.P. performed and analyzed behavioral experiments under the guidance of Y.P. and M.M.; S.O. performed histology and imaging on behavioral mice. R.L. prepared neuronal cultures and viral vectors. M.M. and O.Y. analyzed and interpreted the results and wrote the manuscript.

---

**Supplementary figure 1.**
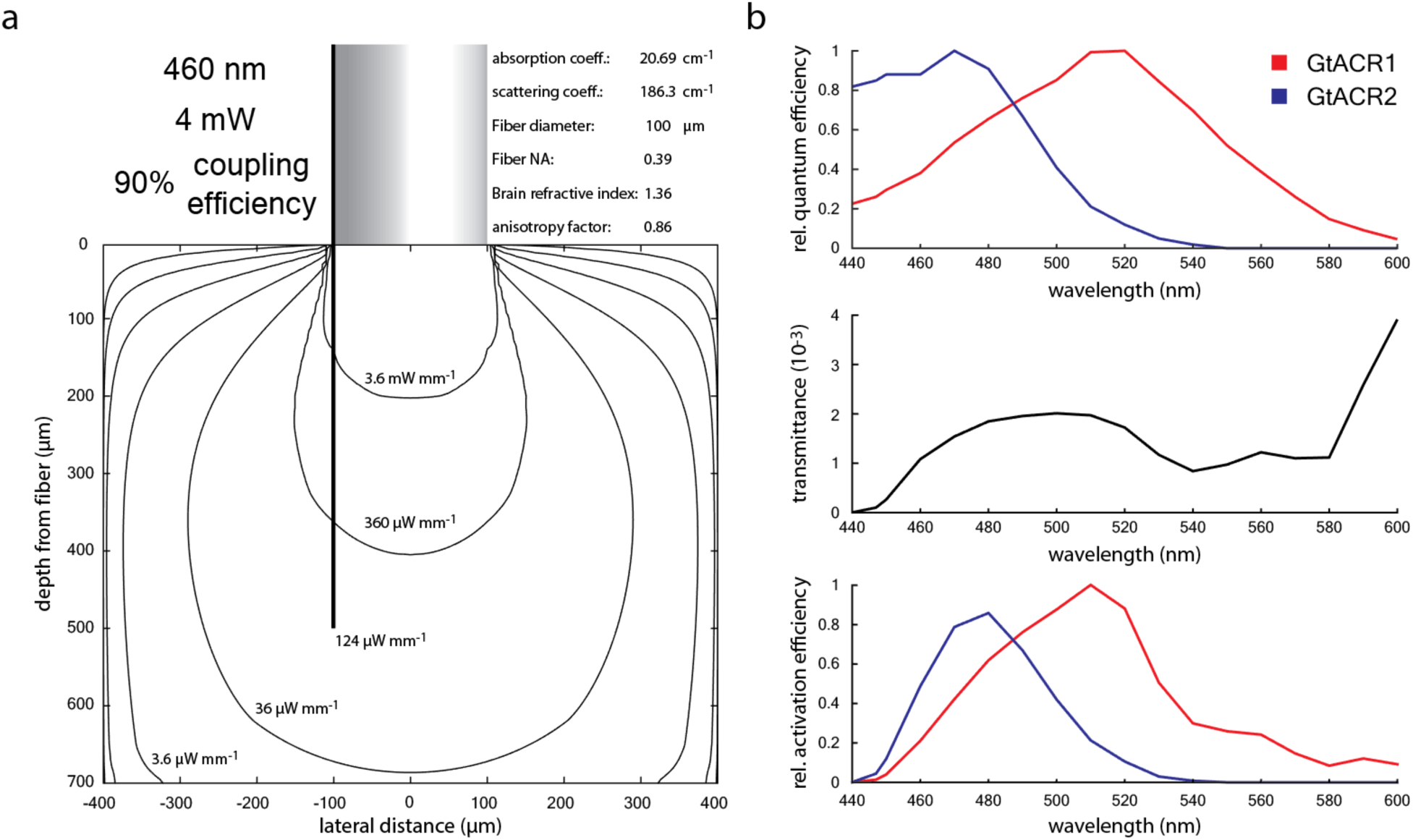
Functional action spectra of GtACR1 and GtACR2 based on multiplication with analytical light spread estimation. Light propagation from a flat-cleaved optical fiber within brain tissue was estimated by analytical modeling ^23^, using gray matter parameters as estimated by Liu et al. ^60^: Wavelength dependent brain scattering coefficiecient: μ_s_(λ) = (2.37*(λ/500nm)-1.15)/(1-g). Absorption is estimated as: μ_a_(λ) = B*S*μ_a_(HbO_2_(λ)) + B*(1-S)*μ_a_(Hb(λ)) + W*μ_a_(H_2_O(λ)). Blood oxygen saturation: S (62%). Estimated percentage of water in brain tissue: W (65%). Percentage of blood in the brain tissue (B). Cerebral cortex (PFC) blood volume excluding major vessels was estimated as 4.6 % according to Chugh et al. (2009) ^61^. Wavelength dependent blood and water absorption coefficients from omlc.org were used. **(a)** Contour lines of estimated light power densities resulting from 4 mW light coupled to an optical fiber. At the recording site (500 μm below and 100 μm lateral to the optical fiber center) the light power density drops to ∼0.11 % of the light power density at the fiber surface. **(b)** Comparison of GtACR1 and GtACR2 activation efficiency within brain tissue by including the wavelength dependent transmittance. Top: relative quantum efficiency as reported in Govorunova et al. (2015) ^16^. Middle: Wavelength dependent transmittance at electrode recording site, modeled as in a. Bottom: Relative activation efficiency normalized by maximal activation of GtACR1. The higher scattering and absorption at 470 nm shift the most efficient excitation wavelength to 480 nm for GtACR2. The higher blood absorption coefficient at 520 nm results in 510 nm being the most efficient wavelength for GtACR1 activation at this distance from the optic fiber. According to this estimation GtACR1 used at 510 nm allows for 14% lower light powers compared to GtACR2 excited at 480 nm, making GtACR1 the more efficient tool when only a single wavelength is needed. However, GtACR1 will cause non-permissive activation in the lower as well as the higher wavelength ranges, therefore only GtACR2 allows for the combination with other currently available optogenetic actuators or reporters.

**Supplementary figure 2.**
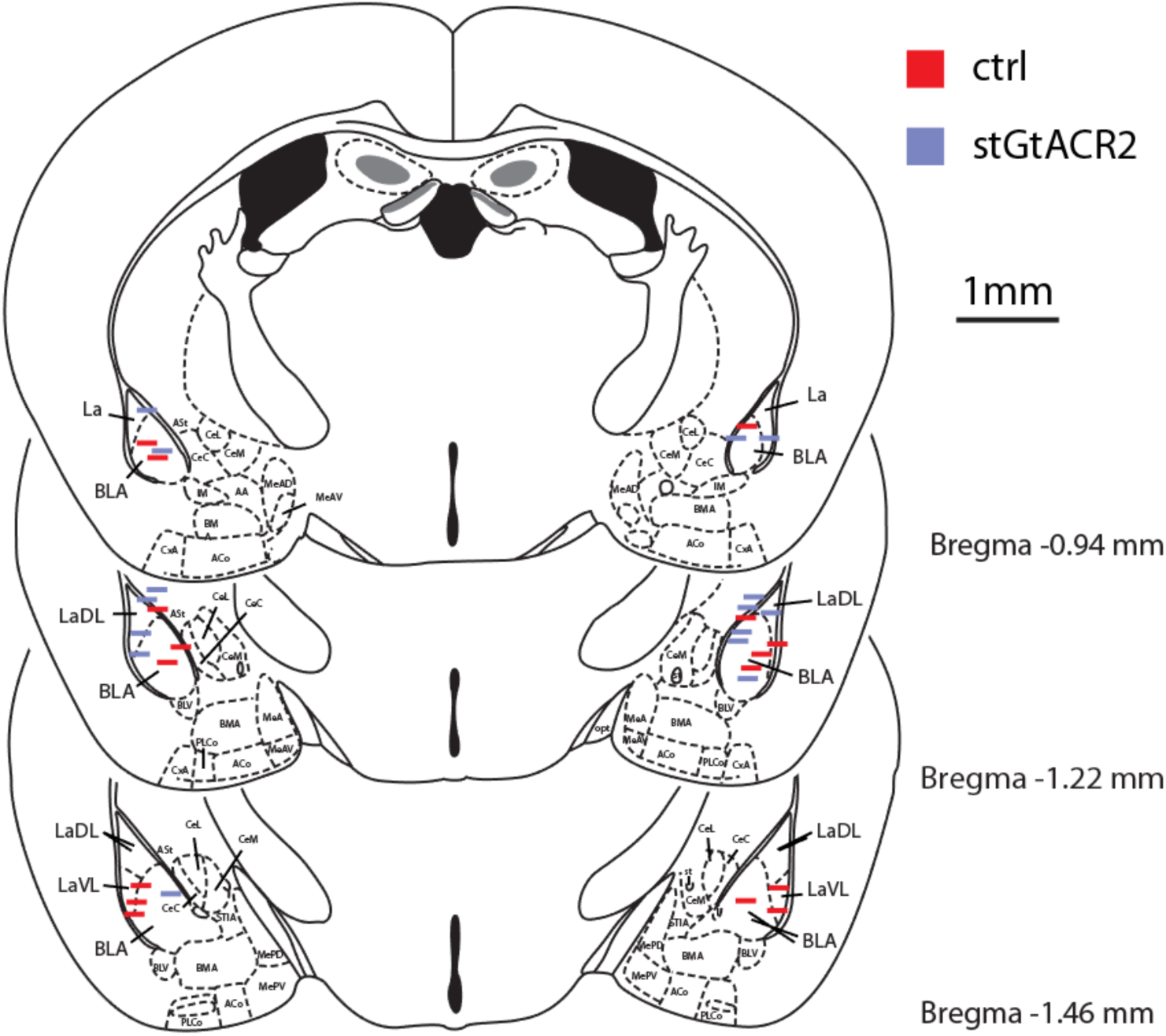
Summary of optical fiber placement of BLA silencing. In fixed brain sections obtained from fear extinction learning experiment animals, fiber placement was determined, and stGtACR2 expression in the BLA close to the fiber was verified. Horizontal lines mark the approximate fiber face position.

## Experimental Procedures

### Production of recombinant AAV vectors

HEK293 cells were seeded at 25%-35% confluence. The cells were transfected 24 h later with plasmids encoding AAV rep, cap and a vector plasmid for the rAAV cassette expressing the relevant DNA using the PEI method ^62^. Cells and medium were harvested 72 h after transfection, pelleted by centrifugation (300 g), resuspended in lysis solution ([mM]: 150 NaCl, 50 Tris-HCl; pH 8.5 with NaOH) and lysed by three freeze-thaw cycles. The crude lysate was treated with 250 U benzonase (Sigma) per 1 ml of lysate at 37°C for 1.5 h to degrade genomic and unpackaged AAV DNA before centrifugation at 3000 g for 15 min to pellet cell debris. The virus particles in the supernatant (crude virus) were purified using heparin-agarose columns, eluted with soluble heparin, washed with phosphate buffered saline (PBS) and concentrated by Amicon columns. Viral suspension was aliquoted and stored at –80°C. Viral titers were measured using real-time PCR. AAV vectors used for intracranial injections had genomic titers ranging between 8.6*10^10^ and 2*10^11^ genome copies per milliliter (gc/ml). Where directly compared virus titers were matched by dilution to the lowest concentration. AAV vectors used for neuronal culture transduction were added 4 days after cell seeding. The titer was matched to final medium concentration of 1.1*10^8^ gc/ml. All of the AAV expression constructs described in this study will be available freely on Addgene to facilitate the utilization of these new tools by the neuroscience community.

The following viruses were used in this study:
AAV2/1.hSyn1.GtACR2-eGFP.WPRE, AAV2/1.CamKIIα.GtACR2-ts-Fred-Kv2.1.WPRE, AAV2/1.CamKIIα.GtACR2-ts-Fred-ER.WPRE, AAV2/1.CamKIIα.GtACR2-ts-Fred-Kv2.1-PEST.WPRE, AAV2/1.CamKIIα.TagRFP-T.WPRE, AAV2/1.CamKIIα.eYFP.WPRE, AAV2/1.CamKIIα.iC++-eYFP.WPRE, AAV2/1.hSyn.iChlOC-eGFP.WPRE.

### Primary hippocampal neuron culture

Primary cultured hippocampal neurons were prepared from male and female P0 Sprague-Dawley rat pups (Envigo). CA1 and CA3 were isolated, digested with 0.4 mg ml^-1^ papain (Worthington), and plated onto glass coverslips pre-coated with 1:30 Matrigel (Corning). Cultured neurons were maintained in a 5% CO_2_ humidified incubator with Neurobasal-A medium (Invitrogen) containing 1.25% fetal bovine serum (FBS, Biological Industries), 4% B-27 supplement (Gibco), 2 mM Glutamax (Gibco) and plated on coverslips in a 24-well plate at a density of 65,000 cells per well. To inhibit glial overgrowth, 200 μM fluorodeoxyuridine (FUDR, Sigma) was added after 4 days of *in vitro* culture (DIV).

### Calcium phosphate transfection of cultured neurons

Neurons were transfected using the calcium phosphate method ^63^. Briefly, the medium of primary hippocampal neurons cultured in a 24 well plate was collected and replaced with 400 μl serum-free MEM medium (ThermoFisher scientific). 30 μl transfection mix (2 μg plasmid DNA and 250 μM CaCl_2_ in HBS at pH 7.05) were added per well. After 1 h incubation the cells were washed 2 times with MEM and the medium was changed back to the collected original medium. Cultured neurons were used between 14 – 17 DIV for experiments.

The following plasmids were used in this study:
pAAV_hSyn1_GtACR2-eGFP_WPRE (based on Addgene 85463), pAAV_ CamKIIα _mNeonGreen_WPRE, pAAV_CamKIIa(0.4kb)_mScarlet_WPRE, pCITF_KCC2-tdTomato (Addgene 61404).

### Animals

All experimental procedures were approved by the Institutional Animal Care and Use Committee (IACUC) at the Weizmann Institute of Science. Six-week-old C57BL/6 mice (P35–45) were obtained from Envigo. Up to 5 male or female C57BL/6 mice were housed in a cage in a light-dark (12 h-12 h) cycle with food and water ad libitum. Mice were housed for 6-12 weeks following surgery to allow for recovery and virus expression.

### Stereotactic injection of viral vectors

Six-week-old C57BL/6 mice (P35–45) were initially induced with ketamine (80 mg kg^-1^) and xylazine (10 mg kg^-1^) and placed into a stereotaxic frame (David Kopf Instruments), before isoflurane anesthesia (^~^1% in O2, v/v). A craniotomy (~1mm in diameter) was made above the injection site. Virus suspensions were slowly injected (100 nl min^-1^) using a 34 G beveled needle (Nanofil syringe, World Precision Instruments). After injection, the needle was left in place for an additional 5 min and then slowly withdrawn. The surgical procedure was either continued with optic fiber or optrode drive implantations (described below), or the surgical incision was closed with tissue glue and 0.05 mg kg^-1^ Buprenorphine was subcutaneously injected for post-surgical analgesia. Injections targeting the medial prefrontal cortex (mPFC) were made 1.8 mm anterior, 0.3 mm lateral and 2.53 mm ventral to bregma. Basolateral amygdala (BLA) injection coordinates were 1.15 mm posterior, 3.0 mm lateral and 5.0 mm ventral to bregma. For mPFC injections, 1 μl of the indicated virus was injected. For fear extinction experiments mice were bilaterally injected with 500 nl AAV2/1.CamKIIα.stGtACR2-Fred.WPRE or AAV2/1.CamKIIα.eYFP.WPRE with a genomic titer in the range of 2-3 × 10^11^ vp ml^-1^.

### Optic fiber and Optrode drive implantation

For fiber optic implantation, a craniotomy (~1 mm in diameter) was made above the implantation site and a ferrule-terminated optical fiber (ThorLabs) was placed at the desired coordinates using a stereotaxic frame (David Kopf Instruments). For bilateral BLA targeting, the fiber tip was placed 1.15 mm posterior, 3.0 mm lateral and 4.8 mm ventral to bregma. For nucleus accumbens, the fiber was implanted at a 45° angle with the ferrule pointing posterior to allow for optrode drive placement above the mPFC in the same animals. The fiber tip was aimed to terminate 1.42 mm anterior, 1 mm lateral and 5 mm ventral to bregma. The optical fiber was secured to the skull using Metabond (Parkell) and dental acrylic. In mice trained for fear extinction learning additional dental acrylic was applied in a second session under isoflurane anesthesia (^~^1% in O_2_, v/v) after fear learning (day 2). For optrode drive implantation, the movable drive was lowered to an initial recording position above the PL (AP: 1.8 mm, ML: 0.3 mm, DV: –2.3 mm). Prior to the permanent attachment of the optrode to the skull, the optrode guide was protected with Kwik-Kast silicone elastomer (World Precision Instruments) and secured using dental acrylic. Mice were allowed to recover for at least 6 weeks before experiments. The locations of implanted optical fibers and optrodes were validated histologically for all experimental mice.

### Acute brain slice preparation

Mice were injected intraperitoneally with pentobarbital (130 mg kg^-1^, i.p.) and perfused with carbogenated (95% O_2_, 5% CO_2_) ice-cold slicing solution ([mM] 2.5 KCl, 11 glucose, 234 sucrose, 26 NaHCO3, 1.25 NaH2PO4, 10 MgSO4, 2 CaCl2; 340 mOsm). After decapitation, 300 μm coronal mPFC slices were prepared in carbogenated ice-cold slicing solution using a vibratome (Leica VT 1200S) and allowed to recover for 20 min at 33°C in carbogenated high-osmolarity artificial cerebrospinal fluid (high-Osm ACSF; [mM] 3.2 KCl, 11.8 glucose, 132 NaCl, 27.9 NaHCO_3_, 1.34 NaH_2_PO_4_, 1.07 MgCl_2_, 2.14 CaCl_2_; 320 mOsm) followed by 40 min incubation at 33°C in carbogenated ACSF ([mM] 3 KCl, 11 glucose, 123 NaCl, 26 NaHCO_3_, 1.25 NaH_2_PO_4_, 1 MgCl_2_, 2 CaCl_2_; 300 mOsm). Subsequently, slices were kept at RT in carbogenated ACSF until use. The recording chamber was perfused with carbogenated ACSF at a rate of 2 ml min^-1^ and maintained at 32°C.

### Electrophysiological methods for cell culture and acute brain slice recordings

Whole-cell patch clamp recordings were performed under visual control using oblique illumination on a two-photon laser scanning microscope (Ultima IV, Bruker) equipped with a 12 bit monochrome CCD camera (QImaging QIClick-R-F-M-12). Borosilicate glass pipettes (Sutter Instrument BF100-58-10) with resistances ranging from 3–7 MΩ were pulled using a laser micropipette puller (Sutter Instrument Model P-2000). For hippocampal neuron cultures, electrophysiological recordings from neurons were obtained in Tyrode’s medium ([mM] 150 NaCl, 4 KCl, 2 MgCl_2_, 2 CaCl_2_, 10 D-glucose, 10 HEPES; 320 mOsm; pH adjusted to 7.35 with NaOH), AcOH Tyrode’s medium ([mM] 125 NaCl, 25 AcOH, 4 KCl, 2 MgCl_2_, 2 CaCl_2_, 10 D-glucose, 10 HEPES; 320 mOsm; pH adjusted to 7.35 with NaOH). The recording chamber was perfused at 0.5 ml min^-1^ and maintained at 29°C. Pipettes were filled using standard intracellular solution ([mM] 135 K-gluconate, 4 KCl, 2 NaCl, 10 HEPES, 4 EGTA, 4 MgATP, 0.3 NaGTP; 280 mOsm kg^-1^; pH adjusted to 7.3 with KOH) or an intracellular solution allowing for EPSC and IPSC recording ([mM] 120 Cs-gluconate, 11 CsCl, 1 MgCl_2_, 1 CaCl_2_, 10 HEPES, 11 EGTA, 5 QX-314; 280 mOsm kg^-1^; pH adjusted to 7.3 with CsOH). Whole-cell voltage clamp recordings were performed using a MultiClamp 700B amplifier, filtered at 8 kHz and digitized at 20 kHz using a Digidata 1440A digitizer (Molecular Devices).

### *In vivo* optical silencing and electrophysiology

All electrophysiological recordings in awake, freely moving mice were performed using an optrode drive consisting of an electrode bundle of 16 microwires (25 μm diameter straightened tungsten wires; Wiretronic Inc.) attached to an 18 pin electrical connector, concentrically arranged around an optical fiber in a mechanically adjustable drive (Anikeeva et al., 2011 ^64^). Extracellular waveform signals were collected using the Digital Lynx integrated hardware and software system (Neuralynx Inc.). The electrical signal was filtered (600–6,000 Hz), amplified using a HS-18-CNR-LED unity-gain head-stage amplifier and digitized at 32 kHz. The electrode–fiber assembly was lowered using the mechanical drive to a new recording site at the end of each recording session, leaving at least 1.5 h before the next session to ensure stable recordings. Optical stimulation was applied through a ferrule-terminated optical fiber (ThorLabs) attached to the patch-chord by a zirconia sleeve (ThorLabs). For optical silencing of mPFC, we used a blue diode laser (λ = 460 nm, Omicron NanoTechnology). Light transmission for each optrode drive was measured with a calibrated power meter (ThorLabs) at the tip of the optical fiber at the end of the experiment. Light power was measured daily before experiments at the tip of the optical patch cord. Neural data were sorted manually using Off-Line Spike Sorter 3.2.4 (OFSS, Plexon) and analyzed in Matlab (MathWorks).

### *In vivo* optogenetic silencing in mice during extinction training

Mice in both the stGtACR2 and control group (eYFP expressing) were placed in the fear conditioning chamber (Med Associates) in context A. Mice were presented with five pairings of the CS (50 ms long 5 kHz 84.4 dB tones, delivered at 10 Hz for 30 s) and US (continuous 0.5 mA foot shock for 1 s). Each CS coterminated with the US, with a 60 s interval between CS–US pairings. On day 3, mice were connected to the optical patch chord and then placed in a different chamber (context B). Context B differed from context A in the following aspects: odor (A: 1 % Acetic acid vs. B: 70 % EtOH), lighting (A: IR vs. B: IR + white light), box size (A: small, B: large), floor texture (A: grid, B: plain), wall texture (A: metal vs. B: Plexiglas), and background noise (A: none vs. B: fan). Mice were allowed 10 min of habituation and then presented with 20 repetitions of the CS, separated by 60 s intervals. The CS was paired with 5 mW blue light (447 nm) administered bilaterally from the fiber tip in both groups. To test extinction learning, mice were placed in context B on day 4 and presented with 20 repetitions of the CS, separated by 60 s intervals. Movies recorded at 25 frames per second were automatically scored for freezing on day 1 and 4 by EthoVision XT 11.5 (Noldus) and by a custom written OpenCV-Python script. The number of changed pixels compared to the last frame was quantified and filtered by a Gaussian filter with 3 frames standard deviation. When mice were connected to optical patch cords, only changed pixels around the mouse body were considered, to discard patch cord motion. A mouse was considered to be freezing if 38 consecutive values (1.5 s) were below 983 pixels (0.5% of all pixels, EthoVision) or 100 pixels (within the ROI around the mouse, OpenCV-Python script).

### Immunofluorescence and microscopy

Hippocampal neuronal cultures were fixed for 15 min with 4% paraformaldehyde in PBS. Coverslips were washed three times in PBS, incubated in blocking solution for 45 min (10% normal donkey serum (NDS) with 0.1% Triton in PBS) and then exposed over night at 4°C to monoclonal mouse anti-KCC2 primary antibody (diluted 1:1500 in 5% NDS, PBS; catalog # 167594 S1-12; USBiological) and rabbit anti-MAP2 (diluted 1:1000 in 5% NDS, PBS; catalog # 4542S; Cell Signaling Technology). Following 3 washes in PBS, coverslips were incubated for 2 h at room temperature (RT) with a Cy5 Donkey Anti-Rabbit IgG (H+L) (diluted 1:500 in 5% NDS, PBS; catalog # 711-175-152; Jackson ImmunoResearch) and Cy3 Donkey Anti-Mouse IgG (H+L) (diluted 1:1000 in 5% NDS, PBS; catalog # 715-165-151; Jackson ImmunoResearch). Coverslips were then washed 2 times with PBS, dipped briefly into double-distilled water and embedded in DABCO mounting medium (Sigma). Immunostained neurons were imaged with a confocal scanning microscope (LSM 700, Carl Zeiss) using a 20 × objective for overview images (NA 0.8; Carl Zeiss) and a 63 × oil immersion objective (NA 1.40; Carl Zeiss) for quantification. Mice were deeply anesthetized using pentobarbital (0.4 mg g^-1^ body weight) and perfused transcardially with ice-cold phosphate buffered saline (PBS, pH 7.4) followed by a solution of 4% paraformaldehyde (PFA) in PBS. After overnight postfixation at 4 °C, brains were removed from the skull and incubated overnight in 4% PFA in PBS. Brains were stored in to 30% sucrose in PBS for at least 24 h or until sectioning. Coronal sections (30 μm or 50 μm) were cut on a microtome (Leica Microsystems) and collected in cryoprotectant solution (25% glycerol, 30% ethylene glycol in PBS pH 6.7). Free-floating sections were mounted on gelatin-coated slides, dehydrated and embedded in DABCO mounting medium (Sigma). Images were acquired using a virtual slide scanner V (Olympus). Acquisition settings were kept constant within each experiment to allow for comparison between mice.

### *In vitro* illumination and drug application

Whole-field illumination *in vitro* was performed using a 470 nm light emitting diode (29 nm bandwidth LED; M470L2-C2; Thorlabs) delivered through the microscope illumination path including a custom dichroic in order to reflect the 470 nm activation wavelength. Light power densities were calculated by measuring the light transmitted through the objective using a power meter (Thorlabs PM100A with S146C sensor) and dividing by the illumination area, calculated from the microscope objective field number and magnification ^65^. D-AP5 (25 μM; ab120003; Abcam) and CNQX (10 μM; C-141, Alomone) were bath applied during all culture experiments. For spatially-restricted illumination of neuronal soma or neurites, a 473 nm diode laser (Bruker) was directed to the imaging plane with galvanometric mirrors, yielding a diffraction-limited spot of light that provided brief light pulses (1 ms) at each location, with 500 ms inter-pulse intervals between non adjacent locations.

### Data analysis and statistical methods

During whole-cell patch-clamp recordings, pClamp 10 software (Molecular Devices) was used for acquisition. Data was analyzed using custom scripts written in Matlab (Mathworks). To quantify postsynaptic current amplitudes in response to light pulses, holding current traces were filtered with a Savitzky-Golay 11 point, second order, Welch window function filter and the maximal change in holding current within 20 ms (EPSCs) after light delivery was determined. Fiji (based on ImageJ2; US National Institutes of Health) was used for immunofluorescence image analysis. In the KCC2 immunofluorescence experiment all numbers (n) refer to the number of imaged axons, in the targeting histology n refers to the number of mice, and in electrophysiological recordings, n refers to the number of recorded neurons / units. To detect significantly modulated units during *in vivo* silencing experiments, a paired-sample student’s t-test was performed comparing the number of detected action potentials between the 5 s pre-light period and the 5 s light period during 10 trials per light power. To detect antidromic spiking units, a paired-sample student’s t-test was performed comparing the number of detected action potentials between the 20 ms pre-light period and the 20 ms light period starting with the 5 ms light pulse. All values are indicated as mean ± SEM. Significance was determined at a significance level of 0.05 with Tukey’s honestly significant difference (HSD) *post hoc* test used to correct for multiple comparisons. In case of non-normal data distribution non-parametric tests were used: Mann–Whitney U test was used for a single comparisons, the Kruskal-Wallis H test for one-way analysis of variance, and the Ray Hare test for two-way analysis of variance. No statistical tests were run to predetermine sample size, but sample sizes were similar to those commonly used in the field. Blinding and randomization were not performed; however automated analysis was used whenever possible.

## References

1. Wurtz, R. H., Using perturbations to identify the brain circuits underlying active vision. Phil. Trans. R. Soc. B 370, 20140205 (2015).

2. Deisseroth, K., Optogenetics. Nature methods 8, 26-29 (2011).

3. Han, X. & Boyden, E. S., Multiple-color optical activation, silencing, and desynchronization of neural activity, with single-spike temporal resolution. PloS one 2, e299 (2007).

4. Zhang, F. et al., Multimodal fast optical interrogation of neural circuitry. Nature 446, 633-639 (2007).

5. Chow, B. Y. et al., High-performance genetically targetable optical neural silencing by light-driven proton pumps. Nature 463, 98-102 (2010).

6. Chuong, A. S. et al., Noninvasive optical inhibition with a red-shifted microbial rhodopsin. Nature neuroscience 17, 1123-1129 (2014).

7. Inoue, K. et al., A light-driven sodium ion pump in marine bacteria. Nature communications 4, 1678 (2013).

8. Mattis, J. et al., Principles for applying optogenetic tools derived from direct comparative analysis of microbial opsins. Nature methods 9 (2), 159-172 (2011).

9. Stujenske, J. M., Spellman, T. & Gordon, J. A., Modeling the Spatiotemporal Dynamics of Light and Heat Propagation for In Vivo Optogenetics. Cell reports 12 (3), 525-534 (2015).

10. Arias-Gil, G., Ohl, F. W., Takagaki, K. & Lippert, M. T., Measurement, modeling, and prediction of temperature rise due to optogenetic brain stimulation. Neurophotonics 3(4), 045007 (2016).

11. Cardin, J. A. et al., Targeted optogenetic stimulation and recording of neurons in vivo using cell-type-specific expression of Channelrhodopsin-2. Nature protocols 5, 247-254 (2010).

12. Groma, G. I. & Dancshazy, Z., How Many M Forms are there in the Bacteriorhodopsin Photocycle? Biophysical journal 50 (2), 357-366 (1986).

13. Mahn, M., Prigge, M., Ron, S., Levy, R. & Yizhar, O., Biophysical constraints of optogenetic inhibition at presynaptic terminals. Nature neuroscience 19 (4), 554-556 (2016).

14. Raimondo, J. V., Kay, L., Ellender, T. J. & Akerman, C. J., Optogenetic silencing strategies differ in their effects on inhibitory synaptic transmission. Nature neuroscience 15, 1102-1104 (2012).

15. Wietek, J. et al., An improved chloride-conducting channelrhodopsin for light-induced inhibition of neuronal activity in vivo. Scientific reports 5, 14807 (2015).

16. Govorunova, E. G., Sineshchekov, O. A., Janz, R., Liu, X. & Spudich, J. L., Natural light-gated anion channels: A family of microbial rhodopsins for advanced optogenetics. Science 349, 647-650 (2015).

17. Berndt, A. et al., Structural foundations of optogenetics: Determinants of channelrhodopsin ion selectivity. Proceedings of the National Academy of Sciences 113, 822-829 (2016).

18. Sineshchekov, O. A., Govorunova, E. G., Li, H. & Spudich, J. L., Gating mechanisms of a natural anion channelrhodopsin. Proceedings of the National Academy of Sciences 112, 14236-14241 (2015).

19. Mohammad, F. et al., Optogenetic inhibition of behavior with anion channelrhodopsins. Nature methods 14 (3), 271-274 (2017).

20. Mauss, A. S., Busch, C. & Borst, A., Optogenetic Neuronal Silencing in Drosophila during Visual Processing. Scientific Reports 7, 13823 (2017).

21. Mohamed, G. A. et al., Optical inhibition of larval zebrafish behaviour with anion channelrhodopsins. BMC biology 15, 103 (2017).

22. Baker, C. A., Elyada, Y. M., Parra, A. & Bolton, M. M., Cellular resolution circuit mapping with temporal-focused excitation of soma-targeted channelrhodopsin. eLife 5 (2016).

23. Yona, G., Meitav, N., Kahn, I. & Shoham, S., Realistic numerical and analytical modeling of light scattering in brain tissue for optogenetic applications. eneuro 3, ENEURO--0059 (2016).

24. Malyshev, A. Y. et al., Chloride conducting light activated channel GtACR2 can produce both cessation of firing and generation of action potentials in cortical neurons in response to light. Neuroscience letters 640, 76-80 (2017).

25. Ganguly, K., Schinder, A. F., Wong, S. T. & Poo, M.-m., GABA itself promotes the developmental switch of neuronal GABAergic responses from excitation to inhibition. Cell 105, 521-532 (2001).

26. Payne, J. A., Stevenson, T. J. & Donaldson, L. F., Molecular characterization of a putative K-Cl cotransporter in rat brain A neuronal-specific isoform. Journal of Biological Chemistry 271, 16245-16252 (1996).

27. Khirug, S. et al., Distinct properties of functional KCC2 expression in immature mouse hippocampal neurons in culture and in acute slices. European Journal of Neuroscience 21, 899-904 (2005).

28. Kelsch, W. et al., Insulin-like growth factor 1 and a cytosolic tyrosine kinase activate chloride outward transport during maturation of hippocampal neurons. The Journal of neuroscience : the official journal of the Society for Neuroscience 21 (21), 8339-8347 (2001).

29. Gabbott, P. L. A., Warner, T. A., Jays, P. R. L., Salway, P. & Busby, S. J., Prefrontal cortex in the rat: Projections to subcortical autonomic, motor, and limbic centers. The Journal of Comparative Neurology 492, 145-177 (2005).

30. Zhu, L., Lovinger, D. & Delpire, E., Cortical neurons lacking KCC2 expression show impaired regulation of intracellular chloride. Journal of neurophysiology 93 (3), 1557-1568 (2005).

31. Hübner, C. A. et al., Disruption of KCC2 reveals an essential role of K-Cl cotransport already in early synaptic inhibition. Neuron 30 (2), 515-524 (2001).

32. Szabadics, J. et al., Excitatory effect of GABAergic axo-axonic cells in cortical microcircuits. Science (New York, N.Y.) 311 (5758), 233-235 (2006).

33. Báldi, R., Varga, C. & Tamás, G., Differential distribution of KCC2 along the axo-somato-dendritic axis of hippocampal principal cells. The European journal of neuroscience 32 (8), 1319-1325 (2010).

34. Gradinaru, V., Thompson, K. R. & Deisseroth, K., eNpHR: a Natronomonas halorhodopsin enhanced for optogenetic applications. Brain cell biology 36 (1-4), 129-139 (2008).

35. Gradinaru, V. et al., Molecular and cellular approaches for diversifying and extending optogenetics. Cell 141 (1), 154-165 (2010).

36. Trimmer, J. S., Immunological identification and characterization of a delayed rectifier K+ channel polypeptide in rat brain. Proceedings of the National Academy of Sciences of the United States of America 88 (23), 10764-10768 (1991).

37. Lim, S. T., Antonucci, D. E., Scannevin, R. H. & Trimmer, J. S., A novel targeting signal for proximal clustering of the Kv2.1 K+ channel in hippocampal neurons. Neuron 25 (2), 385-397 (2000).

38. Wu, C., Ivanova, E., Zhang, Y. & Pan, Z.-H., rAAV-mediated subcellular targeting of optogenetic tools in retinal ganglion cells in vivo. PloS one 8 (6), e66332 (2013).

39. Rogers, S., Wells, R. & Rechsteiner, M., Amino acid sequences common to rapidly degraded proteins: the PEST hypothesis. Science (New York, N.Y.) 234 (4774), 364-368 (1986).

40. Tovote, P., Fadok, J. P. & Lüthi, A., Neuronal circuits for fear and anxiety. Nature reviews. Neuroscience 16 (6), 317-331 (2015).

41. Sierra-Mercado, D., Padilla-Coreano, N. & Quirk, G. J., Dissociable roles of prelimbic and infralimbic cortices, ventral hippocampus, and basolateral amygdala in the expression and extinction of conditioned fear. Neuropsychopharmacology : official publication of the American College of Neuropsychopharmacology 36 (2), 529-538 (2011).

42. Wietek, J. et al., Anion-conducting channelrhodopsins with tuned spectra and modified kinetics engineered for optogenetic manipulation of behavior. Scientific Reports 7, 14957 (2017).

43. Alle, H. & Geiger, J. R. P., GABAergic spill-over transmission onto hippocampal mossy fiber boutons. Journal of Neuroscience 27, 942-950 (2007).

44. Ruiz, A., Campanac, E., Scott, R. S., Rusakov, D. A. & Kullmann, D. M., Presynaptic GABAA receptors enhance transmission and LTP induction at hippocampal mossy fiber synapses. Nature neuroscience 13, 431-438 (2010).

45. Pugh, J. R. & Jahr, C. E., Axonal GABAA receptors increase cerebellar granule cell excitability and synaptic activity. The Journal of neuroscience : the official journal of the Society for Neuroscience 31 (2), 565-574 (2011).

46. Turecek, R. & Trussell, L. O., Presynaptic glycine receptors enhance transmitter release at a mammalian central synapse. Nature 411 (6837), 587-590 (2001).

47. Price, G. D. & Trussell, L. O., Estimate of the chloride concentration in a central glutamatergic terminal: a gramicidin perforated-patch study on the calyx of Held. Journal of Neuroscience 26, 11432-11436 (2006).

48. Kaila, K., Price, T. J., Payne, J. A., Puskarjov, M. & Voipio, J., Cation-chloride cotransporters in neuronal development, plasticity and disease. Nature Reviews Neuroscience 15, 637-654 (2014).

49. Liu, X. et al., Optogenetic stimulation of a hippocampal engram activates fear memory recall. Nature (2012).

50. Kim, C. K., Adhikari, A. & Deisseroth, K., Integration of optogenetics with complementary methodologies in systems neuroscience. Nature Reviews Neuroscience 18, 222-235 (2017).

51. Wiegert, J. S., Mahn, M., Prigge, M., Printz, Y. & Yizhar, O., Silencing neurons: tools, applications, and experimental constraints. Neuron 95, 504-529 (2017).

52. Lee, J. H. et al., Global and local fMRI signals driven by neurons defined optogenetically by type and wiring. Nature 465 (7299), 788-792 (2010).

53. Yizhar, O. et al., Neocortical excitation/inhibition balance in information processing and social dysfunction. Nature 477 (7363), 171-178 (2011).

54. Klapoetke, N. C. et al., Independent optical excitation of distinct neural populations. Nature methods 11 (3), 338-346 (2014).

55. Lin, J. Y., Knutsen, P. M., Muller, A., Kleinfeld, D. & Tsien, R. Y., ReaChR: a red-shifted variant of channelrhodopsin enables deep transcranial optogenetic excitation. Nature neuroscience 16 (10), 1499-1508 (2013).

56. Dana, H. et al., Sensitive red protein calcium indicators for imaging neural activity. eLife 5 (2016).

57. Inoue, M. et al., Rational design of a high-affinity, fast, red calcium indicator R-CaMP2. Nature methods 12 (1), 64-70 (2015).

58. Prakash, R. et al., Two-photon optogenetic toolbox for fast inhibition, excitation and bistable modulation. Nature Methods 9, 1171-1179 (2012).

59. Packer, A. M. et al., Two-photon optogenetics of dendritic spines and neural circuits. Nature Methods 9, 1202-1205 (2012).

60. Liu, Y. et al., OptogenSIM: a 3D Monte Carlo simulation platform for light delivery design in optogenetics. Biomedical optics express 6 (12), 4859-4870 (2015).

61. Chugh, B. P. et al., Measurement of cerebral blood volume in mouse brain regions using micro-computed tomography. NeuroImage 47 (4), 1312-1318 (2009).

62. Grimm, D., Kay, M. A. & Kleinschmidt, J. A., Helper virus-free, optically controllable, and two-plasmid-based production of adeno-associated virus vectors of serotypes 1 to 6. Molecular therapy : the journal of the American Society of Gene Therapy 7 (6), 839-850 (2003).

63. Graham, F. L. & Eb, A. J., A new technique for the assay of infectivity of human adenovirus 5 DNA. Virology 52 (2), 456-467 (1973).

64. Anikeeva, P. et al., Optetrode: a multichannel readout for optogenetic control in freely moving mice. Nature neuroscience 15 (1), 163-170 (2011).

65. Grünwald, D., Shenoy, S. M., Burke, S. & Singer, R. H., Calibrating excitation light fluxes for quantitative light microscopy in cell biology. Nature protocols 3 (11), 1809-1814 (2008).

66. Pugh, J. R. & Jahr, C. E., Activation of axonal receptors by GABA spillover increases somatic firing. The Journal of neuroscience : the official journal of the Society for Neuroscience 33 (43), 16924-16929 (2013).

